# SnpHub: an easy-to-set-up web server framework for exploring large-scale genomic variation data in the post-genomic era with applications in wheat

**DOI:** 10.1101/626705

**Authors:** Wenxi Wang, Zihao Wang, Xintong Li, Zhongfu Ni, Zhaorong Hu, Mingming Xin, Huiru Peng, Yingyin Yao, Qixin Sun, Weilong Guo

## Abstract

**Background:** The cost of high-throughput sequencing is rapidly decreasing, allowing researchers to investigate genomic variations across hundreds or even thousands of samples in the post-genomic era. The management and exploration of these large-scale genomic variation data require programming skills. The public genotype querying databases of many species are usually centralized and implemented independently, making them difficult to update with new data over time. Currently, there is a lack of a widely used framework for setting up user-friendly web servers for exploring new genomic variation data in diverse species.

**Results:** Here, we present SnpHub, a Shiny/R-based server framework for retrieving, analysing and visualizing the large-scale genomic variation data that be easily set up on any Linux server. After a pre-building process based on the provided VCF files and genome annotation files, the local server allows users to interactively access SNPs/INDELs and annotation information by locus or gene and for user-defined sample sets through a webpage. Users can freely analyse and visualize genomic variations in heatmaps, phylogenetic trees, haplotype networks, or geographical maps. Sample-specific sequences can be accessed as replaced by SNPs/INDELs.

**Conclusions:** SnpHub can be applied to any species, and we build up a SnpHub portal website for wheat and its progenitors based on published data in present studies. SnpHub and its tutorial are available as http://guoweilong.github.io/SnpHub/.

## Introduction

Competition in the field of high-throughput sequencing greatly contributes to the reduction of sequencing costs. Currently, one thousand dollars is the cost of sequencing approximately 5 human genomes, 1 hexaploid wheat genome, 6 maize genomes or 50 rice genomes at an average depth of 10×. Whole-genome sequencing is commonly used for species with mid-sized genome such as soybean [1] and maize [2]. Genotyping-by-sequencing (GBS) or whole-genome exon-capture sequencing (WEC) technologies are also frequently used for large-genome species, such as wheat [3]. Currently, many wheat genome studies profile genomic variations on a scale of hundreds or thousands of accessions through WEC [4,5] or whole-genome re-sequencing (WGS) [6].

Plant sciences have experienced a dramatic increase in the available genomic variation data due to the assessment of diverse species and plentiful germplasm resources. Beyond investigating the genetic diversity among individuals, large panels of high-quality genomic variation data have provided valuable resources and great opportunities for identifying trait-related genes, designing markers, constructing gene trees, exploring the evolutionary history and assisting design of molecular breeding. Low-depth re-sequencing data from recombinant inbred line (RIL) populations can assist in the identification of quantitative trait loci (QTLs) for traits of interest. Profiling the genomic variation of TILLING populations in crop studies can benefit the exploration of candidate variations that are rare in nature. The re-use of genomic variation data plays an important role in driving current plant science research.

As a routing pipeline, the raw reads obtained in whole-genome sequencing are first aligned to reference genomes. Then, SNPs and small insertions/deletions (INDELs) are called and stored in the standard variation call format (VCF) files [7]. Although there are great numbers of command-line tools for bioinformaticians to manage and process VCF files, these files are usually quite large. The efficient management of the massive accumulated genomic sequencing data and exploration of these large-scale genomic variation data require computational skills, exceeding the abilities of most biologists.

Some public databases are available for querying sample-specific genomic variations, such as the IC4R database for rice studies [8] and MaizeGDB for maize studies [9]. These public databases are usually based on re-sequencing data, that are generated and maintained by large international consortia. The web servers are implemented independently, providing different functions in exploring the genomic variations. With the increasing number of researchers from generating new data worldwide, it has become impossible to maintain a centralized database that is both up-to-date and comprehensive. There is great demand for implementing a universal platform for building distributed or private web servers for genomic data querying.

Several attempts have been made to implement web-application frameworks. SNP-Seek II creates HDF files for storing genotypic data and utilizes Java Spring and ZK frameworks for implementing the web-application architecture [10]. SNP-Seek II mainly supports data retrieval but is mainly designed for rice studies, and maintaining the complex computing structure requires professional technicians. CanvasDB is designed as a local database infrastructure for managing and retrieving the variation data using the MySQL database and supports filtering functionality and variation detection using R functions [11]. Gigwa v2 also imports VCF files in the NoSQL database, providing both analysis and visualization functions [12]. However, because relational databases are designed for table-structured data, systems such as MySQL are not the optimal method for managing complex genomic variation information, and uncompressed genomic variation data usually require a large amount of memory. SNiPlay3 is based on the Galaxy framework and provides a panel analysis that mainly focuses on whole-genome studies [13]. However, with the rapid accumulation of self-organized genomic variation data, there are still gaps in meeting the great requirements for a uniform, user-friendly, powerful web server framework to with fast and efficient access to massive genomic variation data both locally and in a centralized location, to allow biologists to investigate genomic variations without the need for programming skills.

Here, we developed SnpHub as a uniform web server framework that can be easily set up locally and can be applied by researchers for conveniently management of the massive processed VCF files and used to interactively explore the genomic diversity and perform rapidly analyses in their own labs. SnpHub is designed for rapidly accessing SNP/INDEL data from specific regions and specific sample groups, rather than performing whole-genome analysis. This framework is designed to be species independent, to support scalable variation data and to provide resourceful and extendable functions for re-using and re-analysing genomic variation data.

## Methods

### The general SnpHub framework

The SnpHub framework is designed to be installed in the Linux system, utilizing the Shiny/R framework and integrating several widely used bioinformatics command-line software packages and R packages for analysing and processing genotyping data. SnpHub can be efficiently hosted on a modest computing server, with a local computer installed with R-studio. Rather than performing a whole-genome general analysis, SnpHub provides an efficient way to quickly access data in a local region, filter sites and samples, and generate a genotype table as the intermedia data. To enhance the performance of re-using SNP/INDEL data for in-depth exploration, the intermedia genotype table is stored in random-access memory (RAM) and then used for subsequent analyses and visualizations (Figure 1).

**Figure 1.**
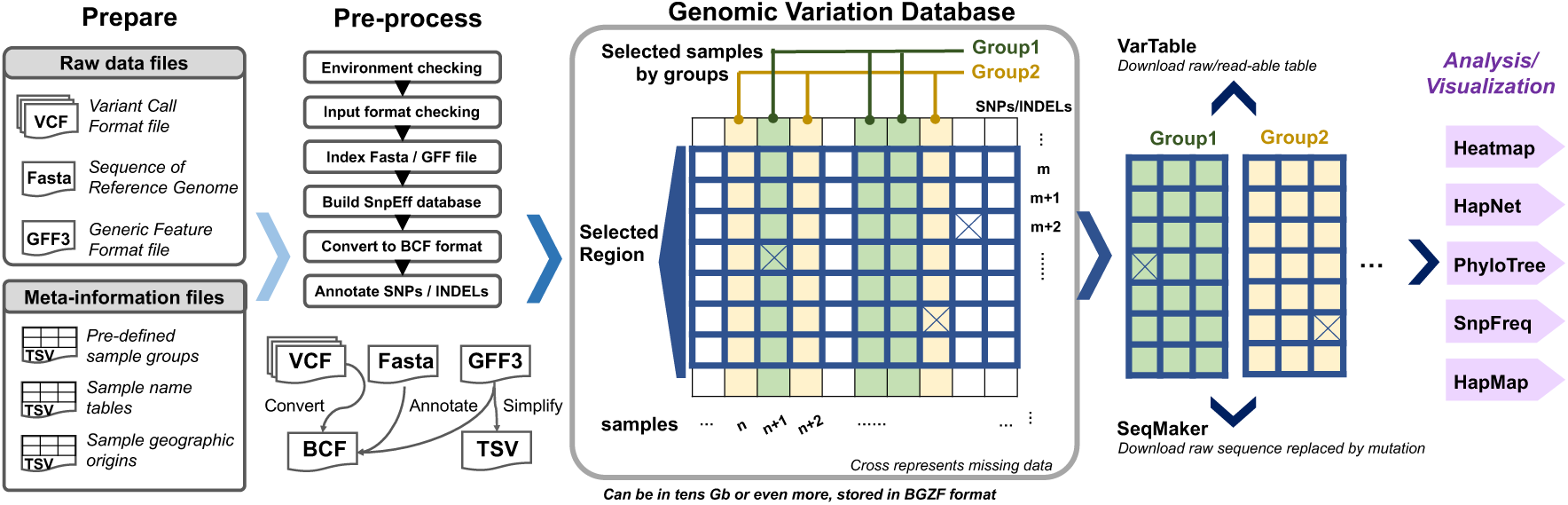
Design schema of the SnpHub server. Once the files and information tables are provided as indicated in the “Prepare” step, the SnpHub server instance performs a pre-processing step for building basic database files and then runs through the Shiny framework. Users can query specific genomic regions or genes for either pre-defined or custom sample groups. SnpHub can efficiently load the raw query data from the hard disk to RAM and then performs an efficient analysis and visualization interactively.

The interactive user interface is implemented using the R/Shiny framework, with powerful, convenient functions for post-processing and visualizing the genotyping data. Considering that a large proportion of open-source bioinformatics analysis tools are implemented using R, SnpHub utilizes the R/Shiny framework to make it compatible and extendable. For simplify installation, a wrapped-version with deploying the SnpHub Docker container is provided at https://github.com/esctrionsit/snphub4docker.

### The setup of SnpHub

#### Prepare step

The SnpHub server-framework is designed to be lightweight and to rapidly access query information from the massive data stored on hard disks while requiring very little RAM. A general Linux workstation (for example, 4G RAM and 2.3 GHz dual process) installed with Shiny/R is enough to set up an instance of SnpHub. Several widely used bioinformatics software programs must be pre-installed, such as SAMtools [14], bcftools [15], seqkit [16] and Tabix [17], along with several R packages.

To build a local instance, the VCF files, reference genome sequence file (FASTA format), gene annotation file (gff3 format) and metadata files defining sample information (tab-separated value, TSV format) are needed. Providing meta-information such as sample information and pre-defined sample groups will enhance performance. A configuration template is provided for presenting meta-information such as the species name, sample description, reference genome, alignment method, and source of the accession.

#### Pre-processing step

A shell wrapper program is provided for the pre-building process. Once an instance server is built, users can access the data through a web browser interactively. Once the configuration information is provided, the local SnpHub instance can be easily built by running the shell-wrapper in one command-line. SnpHub will check the system environment for essential software and the formats of provided files. Then, the gene-based annotation of SNPs/INDELs will be performed by SnpEff [18]. All the meta-information is stored as tables on the hard disks, which is achieved by Tabix for fast retrieval of the content.

### Key features for improving the performance of SnpHub

#### Rapid retrieval of genotype matrix by randomly accessing the disk

Considering that a relational database such as the mySQL framework is suitable for tables and requires a large amount of memory, SnpHub instead adopts the bioinformatician-friendly BCF format for storing the massive genomic variation data. BCF is a binary file format corresponding to VCF [15] with improved performance for supporting the fast querying of a subset of data by randomly accessing the hard disk, taking advantage of the BGZF compression format. In the pre-processing step, all the VCF files will be converted to BCF files. Another benefit is that bioinformaticians can directly perform analysis on these BCF files without storing another copy or format for the same dataset. To improve performance, SnpHub only retrieves a small piece of data for the selected region and selected samples from the disk instantly and stores the intermediate SNP/INDEL table in RAM to be efficiently processed by the downstream analysis functions.

#### The triple-name strategy balances convenience and efficiency

To balance the requirements of convenience in management by server managers, ease of querying and readability of the analysis result, SnpHub utilizes a triple-name/ID for a sample, which includes (a) the *vcfID*, (b) the *Accession name* and (c) the *Display name*. The *vcfID* is a string name that is the same as that provided in the VCF files, avoiding the modification of the original VCF files. The *Accession name* is usually a short name, such as “*S01,S02,S03*”, so that it can be easily typed in the input box for querying a list of samples. The *Display name* is designed as a readable name to be displayed in the results and figures so that researchers can conveniently interpret the result. Arbitrary sample information such as sample passport or sample notes can be provided in additional columns. Once the SnpHub instance is set up, a sample information webpage with a search engine is provided for navigating the names of the available samples and corresponding meta-information.

#### Analysis with defined sample groups

A new feature of SnpHub is that it allows the querying of samples by groups, either using a pre-defined groupID or defining new groups. When setting up the server instance, the database manager can define the system-wide groupIDs by configuring the TSV file. Then, the users can conveniently use groupIDs for querying a list of genes such as *#groupID*. Also, the user can define a custom groupID for a list of samples with the syntax such as *NewGroupA{Sample1,Sample2,Sample3}*. With the defined groups, it will be convenient to refer a list of samples using one GroupID instead of the full list of sample names. By default, SnpHub reserves the group ID “*#ALL*” for querying all the samples in the cohort.

#### Exporting the tables and figures

SnpHub allows users to export tables in CSV format. Beyond interactive visualization of data by the many analysis tools, all the figures can be exported in both PNG and PDF formats. The exported PDF figure represents the vector graphic, as users can conveniently post-edit the figures using tools such as *Illustrator*. A panel of parameters is provided for formulating the height and width of exported figures to produce a satisfactory layout. To be reproducible and traceable, the time-stamp and main parameters are appended to the exported figures.

## Results

### Main functions provided by SnpHub

SnpHub supports the navigation of massive genomic variation data by users by specifying a list of samples and specific genomic regions and performing lightweight analyses and visualizations through the Shiny/R framework. Uniform, flexible interfaces for manipulating the query parameters are provided. As many open-source bioinformatics tools are implemented as command lines or R packages, the Shiny/R framework could be extended for integrating new tools for processing genomic variation tables. SnpHub provides user-friendly functions for navigating genomic variation data by implementing each of the functions on an independent tab page (Figure 2). Raw variation data and genomic sequence retrieval functions are provided in VarTable and SeqMaker. Versatile analysis and visualization functions are provided, including Heatmap, HapNet, PhyloTree, SnpFreq and HapMap. In all of these functions, SnpHub directly queries a gene ID as the corresponding genomic region directly based on the provided GFF file.

**Figure 2.**
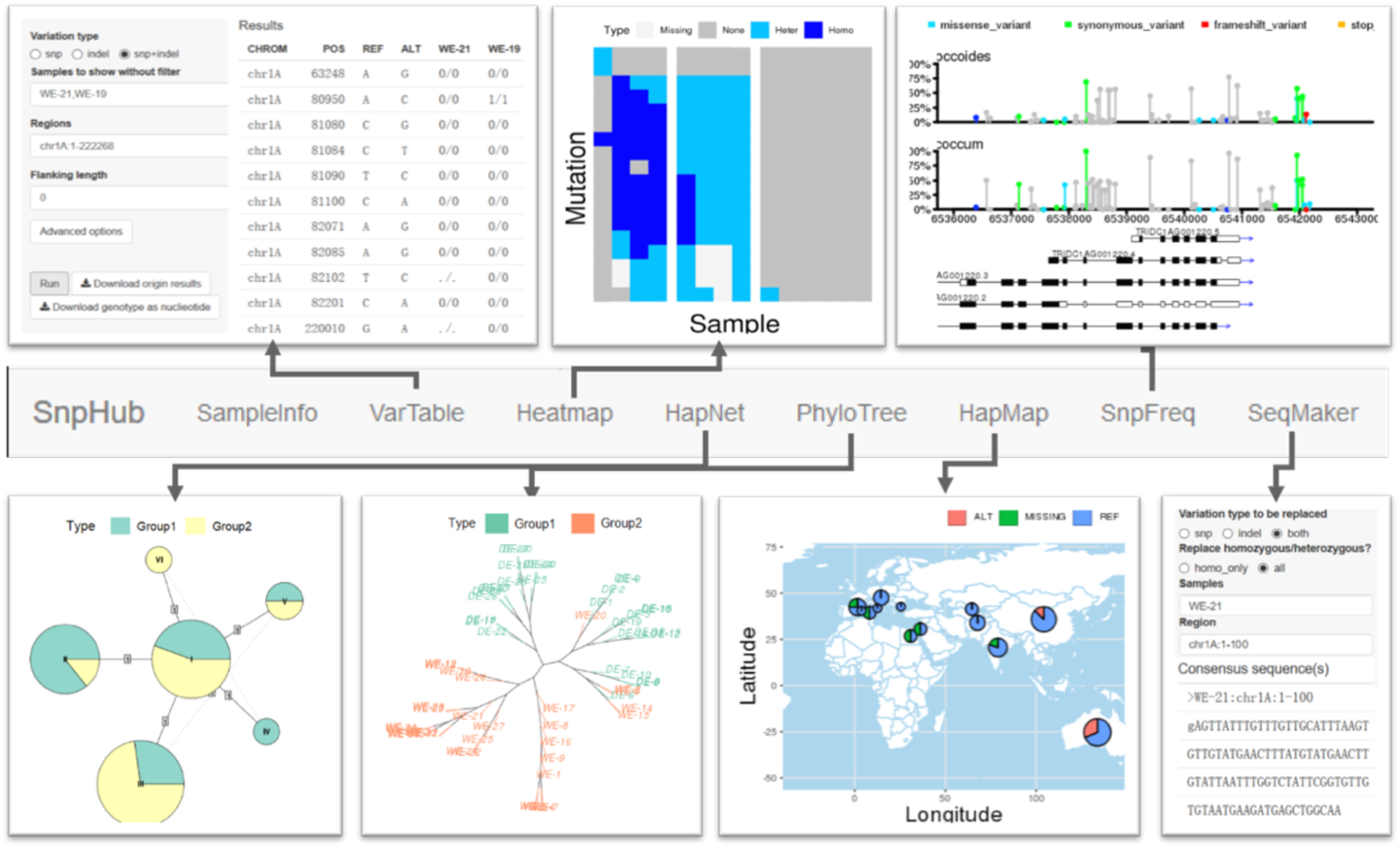
Analysis and visualization functions of the SnpHub server. In one SnpHub instance, each function is implemented in an independent webpage tab.

#### VarTable, for exporting region-specific variation tables

In the ***VarTable*** function, users can query gene-/region-specific SNP/INDEL tables for a list of samples. To be consistent with the VCF format, the exported genotypes are denoted as “0/0”, “1/1”, “0/1” or “./.”, representing the “same genotype with reference genome”, “homozygous variant genotype”, “heterozygous genotype”, or “missing data”, respectively. Tables can be downloaded as raw data or as specific genotypes. This function supports a panel of parameters for filtering sites, such as the minimum allele frequency (MAF) and the maximum missing data frequency. To specify a region, users can either provide a specific region such as “*chr:startPos-endPos*” or use a geneID together with a parameter for the length of the flanking region. To support different purposes for discovering interesting variations, SnpHub extends the sample-based filtering interface to three categories : (a) samples must exhibit genotype variations compared with the reference genome; (b) samples must be consistent with the reference genome in their genotype; and (c) samples shall be exported in parallel without filtering on the basis of genotypes. Beyond the genotype, the meta-data that are stored in VCF format, such as the read depth (DP), genotype quality (GQ) and variant annotations generated by SnpEff (ANN), can also be exported when the optional parameter boxes are checked.

#### Heatmap, for visualizing genotypes in a matrix

The ***Heatmap*** function is an intuitive way to visualize tabular genotype information as a heatmap graph. The samples to be visualized can be provided in one group or a list of groups. By default, genomic positions are displayed along rows, and samples are displayed in columns. The parameters of the two dimensions can be exchanged. Different colours are used to represent homozygous mutations, heterozygous mutations, reference genotypes, and missing data. To be more intuitively visualize possible haplotypes, samples are clustered within each group according to their genotype similarity. This function can be useful for exploring group-specific haplotypes or genotype patterns.

#### HapNet, for constructing a haplotype network

The ***HapNet*** function provides an interface for constructing a haplotype network, which is widely used for characterizing the relationships among population based on sequences. The R package pegas [19] is used for generating the median-joining haplotype network plots. In the HapNet plot, each node represents a haplotype, whose radius is proportional to the number of samples harbouring this haplotype. The distance matrix is calculated among haplotypes based on their sequence distances. Finally, a minimum spanning tree (MST) is constructed. If multiple groups are provided, the nodes will be extended to a pie chart showing the proportion of each group. Similar haplotypes are joined by edges, with the distance shown on the edges. The haplotype network is usually used for exploring the evolutionary paths of different haplotypes among groups of samples [20].

#### PhyloTree, for visualizing sample distance in a local region

The ***PhyloTree*** function supports the exploration of the gene-based genetic distances and evolutionary history based on high-density SNP data. The distance matrix is calculated based on the genetic distances of specified genomic region. Then, two distance-based clustering methods, neighbour-joining (NJ) tree analysis and multidimensional scaling (MDS, also known as PCoA), are supported. NJ-tree analysis cab rapidly evaluate a large amount of data and is suitable for exploring the genetic relationships among samples for a specific region with a low time cost. Versatile layouts for visualizing the NJ-trees are available, including phylogram, cladogram, unrooted, fan, and radial layouts. Samples in different groups are shown in different colours. The MDS analysis supports the visualization of the distances of samples in two-dimensions through non-linear dimensionality reduction. This function provides users with multiple ways to visualize the sample distances for a local region.

#### SnpFreq, for visualizing the SNP annotation in lollipop format

The ***SnpFreq*** function allows users to visualize the SNPs/INDELs and functional annotations along with the transcript-tracks. The previously proposed Lollipop graph [21] is adopted to visualize the positions and frequencies of genomic variations to distinguish the low-frequency variants and un-detected variations. Variants causing amino acid changes are annotated in different colours, including missense variation, synonymous variant, frameshift variant, stop code gained/lost and splice region variants. Transcripts in the same region are displayed as different tracks at the bottom, indicating the exons, introns, CDSs and transcription directions. Samples in different groups are summarized independently and visualized in different tracks, which can be useful for exploring the different frequencies of SNPs between groups.

#### HapMap, for visualizing the genotypes geographically

The ***HapMap*** function provides a way to project the allele distribution of a single site geographically on a map, utilizing the provided resource-gathering locations. A specific genomic site is required for the querying input boxes, such as “*chr:pos*”. If a genomic region is provided, the first variant site in this region will be used for the analysis. To increase user friendliness, this function allows users to adjust the ranges of both longitudes and latitudes. A parameter is provided for the user to select the proper distance for merging geographically closely distributed accessions in one circle. This function could help to shed light on the spreading paths or histories of certain genomic variations/haplotypes.

#### SeqMaker, for creating consensus sequence for an individual

The ***SeqMaker*** function can help to create a consensus sequence by substituting variants based on the reference genome, and the result can be downloaded directly as FASTA file. In principle, this function retrieves a sample-specific sequence by replacing the detected genomic variations, which could be useful for sequence comparisons or primer design. However, it should be noted that the consensus sequences may not reflect the real sequences, considering the missing data as a result of sequencing coverages. Additionally, large structural variants are usually difficult to detect by re-sequencing. By default, “*#RAW*” is preserved for retrieving the raw sequence in the reference genome.

### Construction of the Wheat-SnpHub-Portal by SnpHub

Bread wheat is one of the most important staple crops and exhibits a large, repetitive genome whose genome size is estimated to be ∼16 Gbp. As a hexaploid plant, bread wheat has a complex polyploidization history [22]. Following the release of high-quality reference genomes of wheat and its progenitors, a number of population genomics studies were released together with raw sequencing data or genomic variation data. Jordan et al. sequenced 62 lines of bread wheat (AABBDD) using WEC and GBS methods [23]. Two large WEC-based wheat population genomic studies sequenced 1026 lines [4] and 487 lines [5]. Recently, Cheng et al. performed a high-resolution resequencing study of 93 hexaploidy wheat lines [6]. Population genomics data of wheat progenitors are also available, including data for wild and domesticated emmers (AABB) [24] and of Aegilops tauschii (DD) [25].

We downloaded all the above published datasets (Table 1), and then generated VCF files from raw sequencing data or utilized the published VCF files directly. Thereafter, we constructed up the “*Wheat-SnpHub-Portal*” website, which can be accessed at http://wheat.cau.edu.cn/Wheat_SnpHub_Portal/. Generally, once the configuration data and files are provided, the pre-processing step can be quickly finished, taking from ∼8 minutes [5] to ∼4 hours [6]. Researchers studying wheat or wheat progenitors can access the *Wheat-SnpHub-Portal* and easily explore multiple genomic variation datasets. The *Wheat-SnpHub-Portal* website will be updated with further released genomic variation datasets of wheat and its progenitors in the future.

**Table 1.** The SnpHub instances maintained in Wheat-SnpHub-Portal. WGS, whole-genome resequencing. WEC, Whole-genome Exon-Capture. GBS, Genotyping-By-Sequencing. Disk usage refers to the size of BCF files. *Data is re-analyzed from raw sequencing data.

| Genome | Species | Method | #Sample | Disk usage | Source |
| --- | --- | --- | --- | --- | --- |
| AABBDD | Hexaploid | WGS | 93 | 40Gb | Cheng et al. 2019 |
| AABBDD | Hexaploid | WEC | 487 | 192Mb | Pont et al. 2019 |
| AABBDD | Hexaploid | WEC | 1026 | 1.8Gb | He et al. 2019 |
| AABBDD | Hexaploid | WEC&GBS | 62 | 2.4Gb* | Jordan et al. 2015 |
| AABB | Tetraploid | WEC | 64 | 645Mb | Avni et al. 2017 |
| DD | Diploid | GBS | 567 | 234Mb* | Singh et al. 2019 |

## Discussions

With the decreasing sequencing costs, increased numbers of samples and species will be sequenced. T will be difficult for universal and centralized databases to satisfy the versatile needs for variant analysis and querying new datasets. SnpHub can be applied to any species with an assembled genome and gene annotations. It can be instantly set up based on the VCF files. For the future population genetic studies, a SnpHub querying server ca be easily set up in addition to the publication of the raw data generated, making the data to be more easily accessible by the community. SnpHub can serve as laboratory-level web server for navigating and visualizing the genomic diversity or individual line or lineage. SnpHub can be useful for different occasions: investigators can infer trait-associated genes with population structure information and variation function annotations from specific sample sets; and breeders can access the genetic diversity at specific loci for designing new breeds. SnpHub provides a uniform server framework for easily setting up distributed servers for genotyping-queries and analysis, and can be used to build database portals such as our *Wheat-SnpHub-Portal*, extending this strategy from wheat to other important crops or other plants.

## Availability of supporting source code and requirements

Project name: SnpHub

Project home page: https://guoweilong.github.io/SnpHub/

Operating system(s): Linux

Programming language: R, Shell

Other requirements: R/Shiny, samtools, bcftools, seqkit, tabix

License: MIT licence

## Authors’ contributions

Method development: W.W., Z.W., W.G.; implementation: W.W., Z.W., X.L., W.G.; data preparation: Z.W.; design and testing: W.W., Z.W., Z.N., M.X., H.P., Y.Y., Q.S., W.G.; definition of research project: W.G.

## Acknowledgements

We thank Xiaoming Xie, Yongming Chen, Zhengzhao Yang for exploring technology, and thank Kuohai Yu for IT support.

